# Gum Odina, a novel prebiotic, promotes the expression of sealing tight junction proteins in colon organoids developed from the C57BL/6 mice model

**DOI:** 10.64898/2026.02.03.703214

**Authors:** Sohini Sikdar, Paulami Dutta, Debmalya Mitra, Amalesh Samanta

**Author notes:** Corresponding author: Debmalya Mitra, Associate Research Scientist, Columbia University Irving Medical Center, 630 West 168th Street, New York, NY 10032, United States. **Disclosures:** The authors report no competing interests.

## Abstract

Maintaining gut-microbiome homeostasis is the biggest issue worldwide as per public health concerns. Gut probiotics not only inhibit pathogen invasion in the systemic circulation but also help us metabolize complex food. Therefore, for decades, gut dysbiosis has been proven to be the gateway to several diseases, leading to comorbidity and even mortality. Prebiotics are natural products, mainly nondigestible food ingredients, that help the selective growth of probiotic bacteria in the gut. This study focuses on the novel Gum-Odina (GO) prebiotic and its efficacy on gut microbial metabolite modulation and maintaining gut barrier integrity. Gut wall enterocytes are integrated by a series of tight junctional (TJ) proteins. This study explains the effect of GO prebiotic-modulated gut metabolites on tight-junctional (TJ) protein expression in a murine colon Organoid model. Fecal microbiota from a colitis patient were used to inoculate the SHIME gut simulator, comprising a colitis control run and a Gum-Odina-supplemented run to enrich commensal bacteria selectively. Metabolites from both groups were then applied to healthy colon organoids. According to the mRNA expression analysis, tight junctional sealing proteins such as Zonula occludens, or ZO-1, Occludin, Claudin-1, 4, and 5 were significantly upregulated in the colon organoids upon Gum-Odina administration, whereas no change in the Junctional Adhesion Molecule-A or JAM-A was observed. Downregulation of sealing TJ proteins is the Hallmark of Leaky gut, which was successfully reversed using the Gum-Odina supplement. Hence, Gum-Odina prebiotics have a promising capability to reduce colitis-induced gut permeability and can be considered to be a therapeutic agent in the future.

## Introduction

The organoids represent a three-dimensional (3D) cell culture model which is generated i*n vitro* and can recapitulate the major properties of primary cells, such as cell behaviour, and its function ^**[1]**^. These are organs grown in the lab from adult cells that resemble their parent organ, making them very useful in drug testing and studying disease states at an individual bjklevel. Organoids overcome the limitations caused by both 2D cell culture, which often fails to express all the physiological properties of primary cells, and in-vivo animal models, which are often criticized for their inability to precisely mimic human diseases and reproducibility ^**[2]**^.

Colitis refers to the inflammation of the colon, which can often have an autoimmune or infectious predisposition. Previously, a low incidence of inflammatory bowel diseases (IBD) was reported in Asia, ^**[3,4]**^ however enthralling evidence from other studies has demonstrated a steep rise in the incidence of IBD in regions across Asia ^**[5-8]**^. Poor dietary patterns and stress have accelerated the incidence of colitis with a worldwide estimate of 6.8 million cases ^**[9, 10]**^. The exact cause of colitis is yet to be explored as it is a multifaceted disease but it is clear that colitis involves gut dysbiosis which causes an imbalance between various inflammatory markers. The pathogenic bacteria replace the commensal organisms, and disrupt the gut barrier integrity leading to epithelial damage and abrupt immune responses ^**[11]**^.

With the rapid rise in the incidence of non-communicable diseases of the gut, it is important to reduce the burden of colitis with a newer approach of using biotherapeutics like prebiotics. Prebiotics are non-digestible complex carbohydrates that can modulate the gut microbiota by stimulating the growth of beneficial bacteria in the colon and reducing chronic disease progression ^**[11]**^. Gum Odina (GO), is an established prebiotic which prevents colitis progression in a Dextran Sodium Sulfate (DSS) induced colitis mouse model. GO is derived from *Odina wodier* plant and is a polysaccharide comprising galactose and arabinose as major constituents followed by uronic acids which are accountable for antioxidant properties. GO is fibrous in nature, absorbing deleterious substances from the gut environment and increasing the bulk of feces. GO also liberates short-chain fatty acids (SCFAs) upon degradation, which promotes bacterial diversities when tested on a human gut simulator, thereby reducing gut dysbiosis. The anti-inflammatory properties of GO have already been tested in the colitis mice model ^**[12, 13]**^.

The gut organoids have been a promising tool in evaluating the gut barrier integrity. As the disruption of the gut barrier is one of the major phenomena of colitis, we used DSS ∼50 KD as a chemical colitogen in the culture model to mimic the disease state. Thus, this study explores the effect of GO metabolites on the expression of tight junctional proteins in gut organoids developed from the intestinal cells of C57BL/6 mice and compares the result of disease prognosis and reproducibility with that of the DSS induced colitis mice model.

Moreover, this study will further highlight the potential of working in organoids developed from human tissues, which will enable us to mimic the disease as well as its prognosis at an individual level and it will act as a bridge for future clinical trials.

## Methodology

### Preparation of test metabolites for organoid culture using Simulator of Human Intestinal Microbial Ecosystem (SHIME)

SHIME is an *in vitro* gut model mimicking the human gastrointestinal tract and physiological conditions. The nutritional medium used in SHIME is detailed in Table 1. The start-up and adaptation period was carried out for 2 weeks. The bacteria used for inoculation were a faecal sample of a male colitis patient collected from ID & BG hospital as a specimen abiding by the Institutional Ethics Committee of ICMR NICED, Kolkata vide Ref no.: (A-1/2015-IEC). The test period was conducted for 2 weeks; during this time, samples were collected from the 3rd, 4th, and 5th compartments of SHIME. The samples collected were filtered through 0.45μm and 0.22μm filters and kept at -20 C, which were used as colitis metabolites (C-M) in Organoid experiments. A similar SHIME run was done during the prebiotic phase with the same stool sample, where sucrose in the SHIME media was substituted with GO, and the metabolites collected were named as GO metabolites (GO-M).

### Culturing of colon organoids

From a 8 weeks old male control C57BL/6 mouse, 3-4 cm of the colon was harvested from just below the cecum. Colon pieces were sectioned into 2mm lengths, further cut open longitudinally, and washed 15-16 times with cold PBS. The tissue pieces were transferred into a gentle cell dissociation reagent at 25 °C for 40 min on a rocker. The fragments were taken into a centrifuge tube, allowing them to settle at the bottom. The supernatant was discarded, and the colon pieces were washed with PBS containing 0.1% BSA and passed through a 70 μm cell strainer. The filtrate is collected and labelled as fraction 1. Similarly, fractions 2-5 were prepared by repeating the washing to obtain a fraction that is devoid of debris and fibrous tissue. The tissue fragments were centrifuged at 290 g for 5 min at 2-8 °C. The supernatants were discarded, and the pellet was resuspended in a 15 ml centrifuge tube containing fresh cold PBS buffer with 0.1% BSA and centrifuged again. The fractions suitable for organoid cultures were carefully selected under observation under a microscope and mixed in cold DMEM/F-12. 150 μl of each complete organoid medium (#06005, IntestiCult™ Organoid Growth Medium, Mouse, STEMCELL Technologies) with gentamycin (50 μg/ml) and Matrigel Matrix was added to the tubes containing crypts and mixed gently with a pipette. 50 μl of this mixture is plated on 24-well plates, forming domes. The plate was incubated at 37 °C for 10 min, and 750 μl of complete media was added to wells containing organoids and Matrigel matrix domes. The unused wells are filled with PBS and incubated at 37 °C in a 5% CO2 incubator. Mature organoids were divided into three groups: (a) Control, (b) C-M, and (c) C-GO-M. The wells marked with Colitis and GO were supplemented with 10% Colitis metabolites, and the wells belonging to the GO group were supplemented with 10% GO metabolites on the 3^rd^, 5^th,^ and 6^th^ day. A time-dependent growth of the groups was documented and compared during this experiment.

### Quantification of mRNA expression of tight junction proteins

On day 7^th^ of the experiment, the Organoids were harvested from Matrigel domes for RNA isolation by gently aspirating the culture medium and adding 300 µL Gentle Cell Dissociation Reagent (Catalog #07174, STEMCELL Technologies) to the exposed dome for 5 minutes at room temperature. The domes were disrupted by gentle pipetting ∼10 times and collected in microcentrifuge tubes. The tissue fragments were allowed to settle by gravity for 30 seconds before removing the supernatant. The organoid pellets were washed twice with 1X PBS and then allowed to settle at the bottom for approximately 30 seconds. 500 µL of TRIzol reagent (Invitrogen, Massachusetts, USA) was added to each organoid group for RNA extraction and quantified using nanodrop (NanoDrop 8000 Spectrophotometer, Thermo Fisher Scientific, USA). 5 μg RNA was taken for reverse transcription with specific primers. dNTPs, MuLV reverse transcriptase, and other reagents were added according to the kit manual (Ambion INC, Austin, USA) to produce cDNA of GAPDH, Claudin 1 (*Cldn -1*), Claudin-2 (*Cldn-2*), Claudin-4 (*Cldn-4*), Claudin-5 (*Cldn-5*), Occludin (*Ocln*), Zonula Occludens (*Tjp1*), Junctional adhesion molecule A (JAM-A), and GAPDH was taken as a loading control.qPCR was conducted using SYBR green mix (Qiagen, Maryland, USA) as per the provided protocol. The primers used in the study are detailed below.

*Cldn -1* (>NM_016674.4) (5□ATGCAAAGATGTTTTGCCACAG3□) (Tm=58.60); (5□TACAAATTCCCATTGCAGCCC3□) (Tm=59.17), product length= 210

*Cldn-2* (>NM_001410421.1) (5□TATGTTGGTGCCAGCATTGT3□) (Tm=58.08); (5□TCATGCCCACCACAGAGATA3□) (Tm - 58.12), product length = 205

*Cldn-4* (>NM_009903.2) (5□TCGTGGGTGCTCTGGGGATGCTT3□) (Tm=65.0);

(5□GCGGATGACGTTGTGAGCGGTC3□) (Tm=62.8), product length = 170

*Cldn-5* (>NM_013805.4) (5□-CTTTGTTACCTTGACCGGCG-3□) (Tm=59.48); (5□-CCCTGCTCGTACTTCTGTGA-3□) (Tm=59.11), product length = 198

*Ocln* (>NM_001360538.1) (5□TCACTTTTCCTGCGGTGACT3□) (Tm=59.53); (5□GGGAACGTGGCCGATATAATG3□) (Tm - 58.93), product length = 138

*Tjp1* (>NM_001417368.1) (5□-TCTTCCATCATTTCGCTGTGT-3□) (Tm= 57.94); (5‘-TCTGAAACCATCAAGTCCACA-3’) (Tm= 57.14), product length = 107

mJAM-A (>NM_172647.2) (5□CTGATCTTTGACCCCGTGAC3□) (Tm=58.27); (5□ACCAGACGCCAAAAATCAAG3□) (Tm=56.9), product length = 187

*Gapdh* (>NM_001411843.1) (5□ACCCAGAAGACTGTGGATGG3□) (Tm=59.01); (5□CACATTGGGGTAGGAACAC3□) (Tm - 55.49), product length = 170

### Immunofluorescence imaging for tight junctional protein expression analysis

For confocal imaging, mouse colon organoids were cultured within the removable, 0.4cm^2^ 16-welled chambered microscope slides (Lab-Tek Glass Chamber Slide 16 Well PK96) and grouped into (i) Control, (ii) C-M, (iii) C-GO-M, and (iv) LPS. LPS, at a concentration of 100 μg/mL, was used to induce barrier damage ^**[14]**^. After removing the growth media, organoid cultures were gently rinsed twice with 1× phosphate-buffered saline (PBS; STEMCELL Technologies, Cat. No. 10010). Matrigel domes containing organoids were preserved for 30 minutes at room temperature using 100 µL of 4% paraformaldehyde (PFA). Afterward, the PFA was aspirated slowly and very carefully from the sides of each well, and the Matrigel domes containing organoids were washed twice with 1X PBS. Samples were permeabilized and blocked using a blocking buffer composed of 5% horse serum and 0.5% Triton X-100 prepared in 1× PBS for 2 hours at room temperature. After diluting primary antibodies in blocking buffer (100 µL per well), the organoids were incubated at 4 °C overnight. The samples were cleaned three times using 1× PBS. The secondary antibodies were diluted in blocking solution, applied, and incubated at 4 °C for 2 hours before three further PBS washes. For nuclear staining, samples were incubated at room temperature for 10 minutes with Hoechst. A Carl Zeiss microscope was used to image the mounted samples following three final PBS washes. All the images were processed and analyzed using ZEN.

### Statistical analysis

The experiments were conducted in triplicate in triplicate and data were expressed as mean ± standard error of the mean (SEM), one-way analysis of variance (ANOVA) was conducted followed by Tukey’s multiple comparison tests using Graph Pad Prism 6 software (Graph Pad Software Inc., San Deigo, CA, USA) and difference with P < 0.05 was considered significant (Mitra et al. 2017) ^**[15]**^.

## Results

### Growth of organoids

Figure 1-A represents the sequential bright-field images of organoids, illustrating the growth kinetics from an initial crypt to mature structural complexities. Till day 2 or 3, most of the colon organoids appeared as small spherical structures with a compact mass of cells. Which gradually increased in size, forming well-defined cystic organoids with a distinct outer epithelial boundary on days 3 to 4. Lumen development and increased cell proliferation and complexity are seen in later phases from day 5 to 10, which also include multilobulated formations, asymmetric development, and thickening of the epithelium. These characteristics show that mature colon organoids typically form crypt-like domains. On day 10 and onwards, the organoids naturally started to disintegrate. Figure-1-B represents the recorded growth of Control, Colitis metabolite (C-M), and GO metabolite-supplemented (C-GO-M) wells on day 3, 5, and 7. At first, the number of mature spheroids per microscopic field was higher in the control well compared to the C-M and C-GO-M wells. From the images, it was observed that the integrity of the colonic organoids in the control group was intact, with cystic structures consisting of a hollow lumen surrounded by columnar epithelium, which also exhibited an early budding stage at day 5. On the other hand, the C-M-treated organoids exhibited structural disintegration and loosening of cells from the middle and outer epithelial layers. It also represents that the C-M group had compact masses of cells without lumens. Further, upon treatment with GO metabolites, the growth pattern of the organoid in the C-GO-M group tended to have similar structural integrity with spherical architecture, forming a distinct lumen and cohesive epithelial layer to the control wells. On day 7, the control organoids exhibit a well-morphed, highly folded epithelial layer, indicating the crypt and villus architecture; interestingly, the C-GO-M organoids slowly achieved an early mature form, consisting of some smaller outer budding of the epithelial layer with a narrow lumen in the middle. However, the C-M organoids showed a completely deformed architecture, giving the appearance of being broken up or collapsed. The cellular debris and irregular mass development depict a gradual decrease in the organoid viability and organization over time.

**Figure 1-A.**
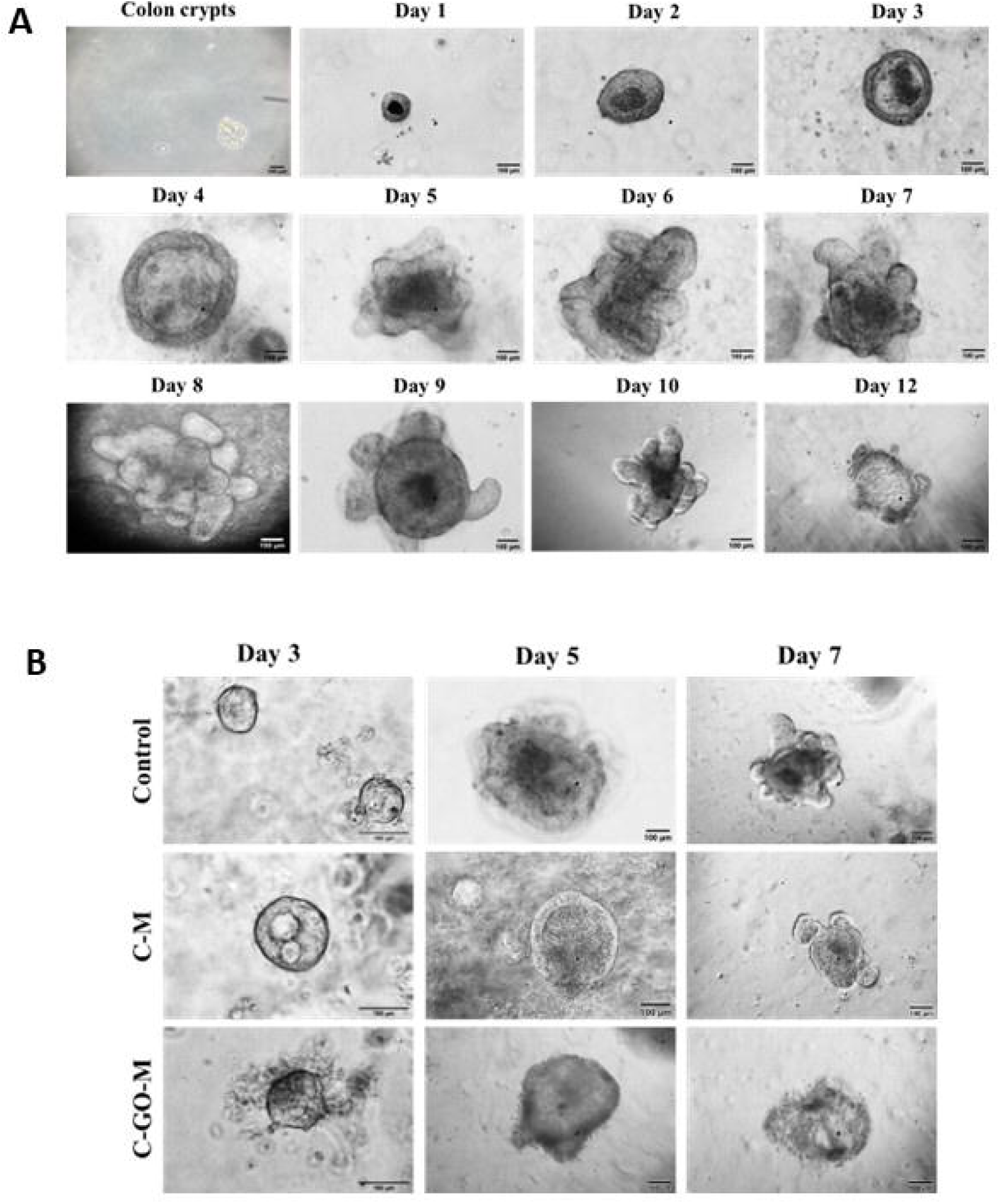
Day-wise growth of colon organoids from colon crypts isolated from C57BL/6 mice. Images were taken under a compound light microscope with the 100µm scale bar (bottom left corner). **B**. Comparative growth rate and growth pattern of control, GO-M, and C-GO-M treated organoids.

### Expression of tight junctional proteins

From Figure-2 it was observed that the relative expression of the sealing tight junction proteins like *Cldn-1, Cldn -4, Cldn-5, Ocln* and *Tjp1* were 0.030 ± 3.47, 0.023 ± 1.83, 0.006 ± 3.27, 0.066 ± 0.95, 0.032 ± 3.38 which got significantly (p ≤ 0.05) reduced by 93.55%, 63.63%, 83.12%, 66.15%, 73.30% respectively in C-M grouped wells with respect to control. After the introduction of GO metabolites in the C-GO-M grouped wells, the expression of *Cldn-1, Cldn-4, Cldn-5* and *Ocln* significantly (p ≤ 0.05) normalised by 51.85%, 71.15%, 50.32% and 50.76% respectively when compared to control. No notable change was observed in the expression ZO-1 upon treatment with GO metabolites. There was a major decrease in the expression of Jam-A in the colitis-grouped wells which increased in GO-grouped wells. The expression of pore-forming protein *Cldn-2* was 0.024 ± 4.87 which got increased to 0.039 ± 4.65 in colitis-grouped wells but we did not see any remarkable change or downregulation in the *Cldn-2* in the GO metabolite treatment. Additionally, the immunofluorescence assay to further investigate the protein expression and distribution of the tight junction protein in the organoids justified the qPCR data by showing similarity.

**Figure 2-.**
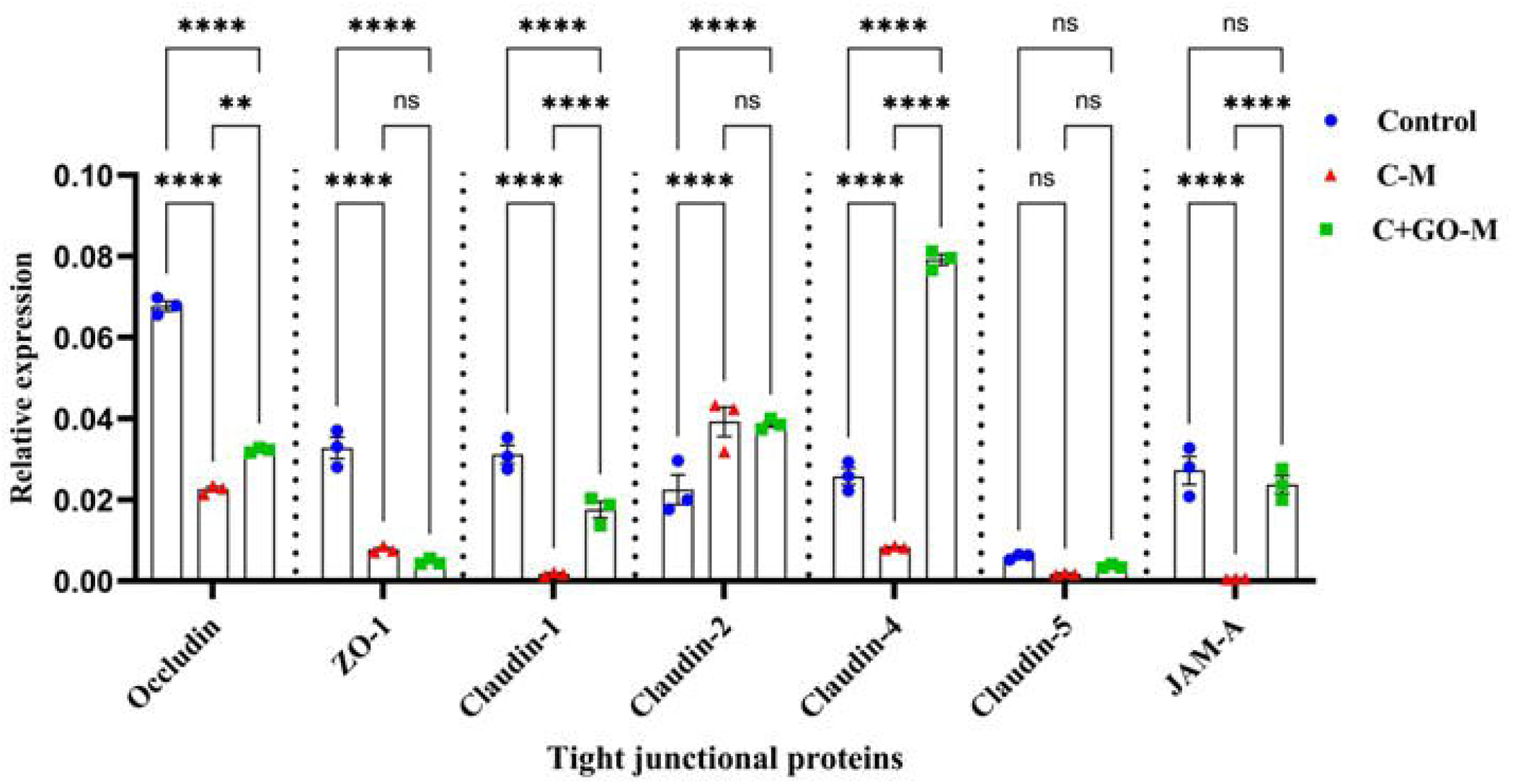
Tight junction expression analysis by real-time PCR. Relative expression change of Occludin, ZO-1, Claudin-1, Claudin-2, Claudin-4, Claudin-5, and JAM-A among Control, G-M, and C-GO-M treated organoids was analyzed.

**Figure 3A-B-.**
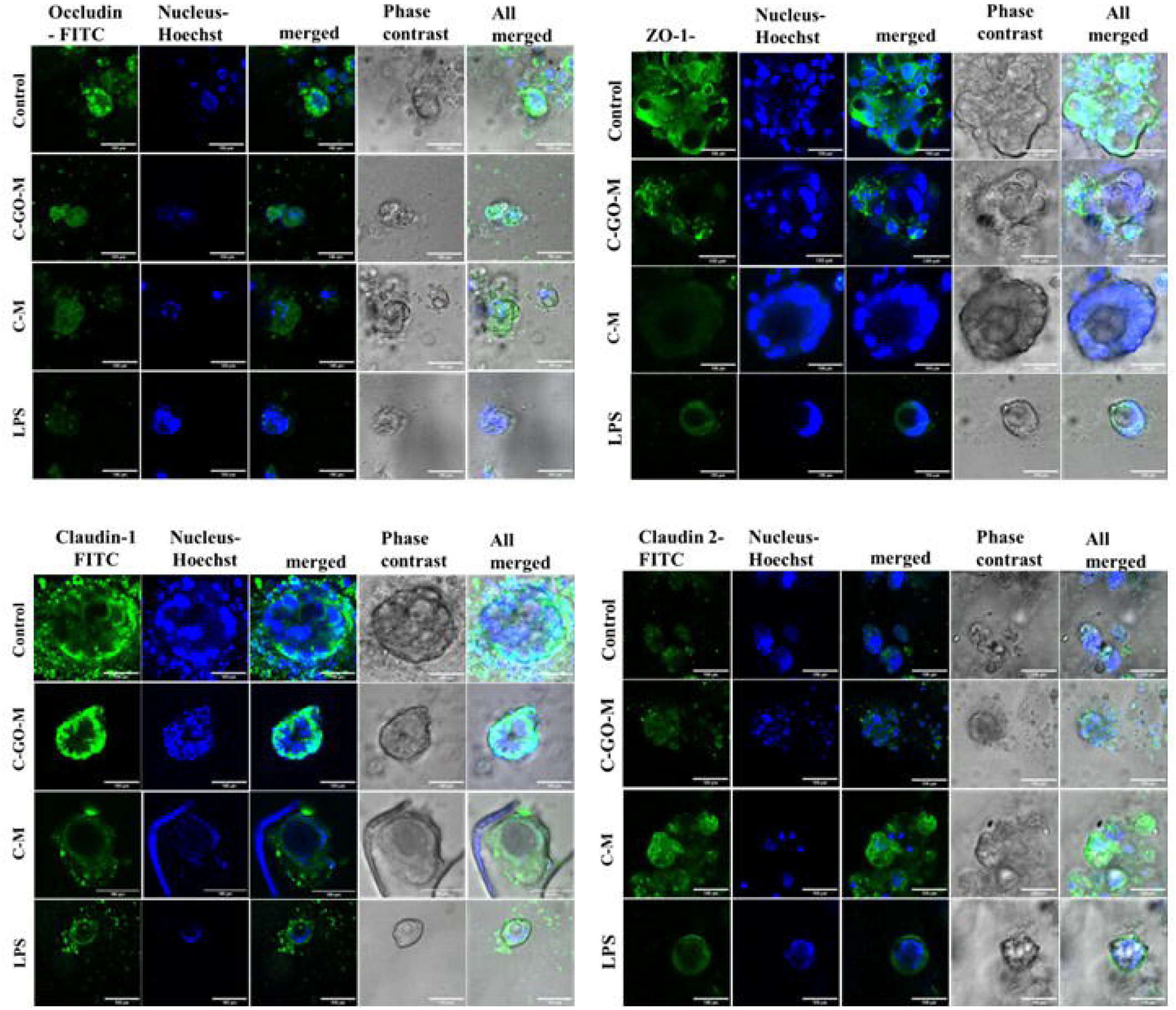
Tight junction expression analysis by FITC (green) tagged anti-Occludin, anti-ZO-1, anti-Claudin-2, anti-E cadherin, anti-Claudin-4, Claudin-5, anti-JAM-A, and Rhodamine (red) tagged anti-F actin. Hoechst (blue) was used as the nuclear stain. Images were taken using a confocal microscope with a 100 μm scale bar.

**Figure 3B.**
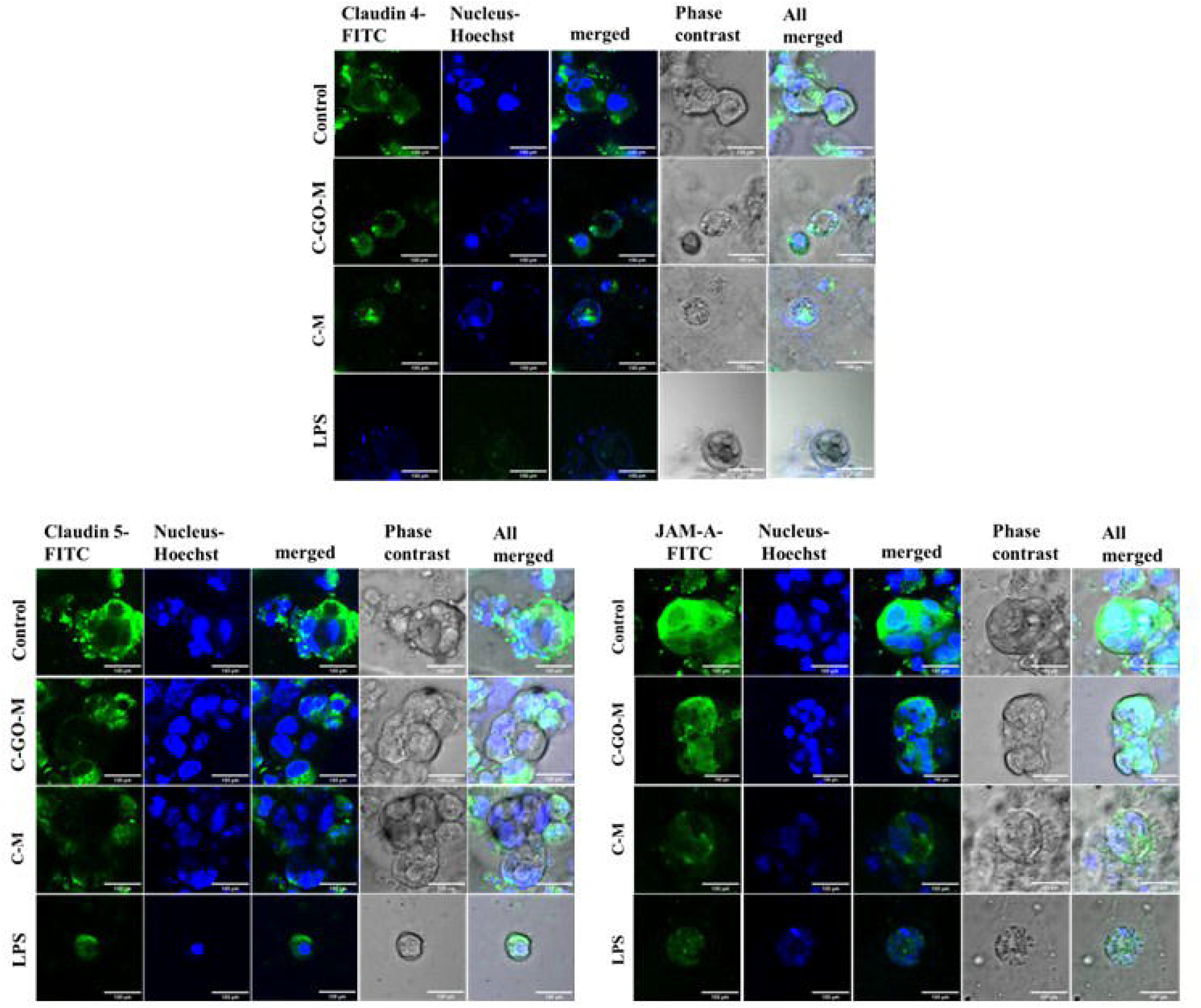

## Discussion

Organoid culture has emerged as a significant translational model in research due to its ability to mimic the structure and function of an organ while preserving the heterogeneity in a laboratory. It has become very efficient in disease modelling and drug testing compared to the single-cell type culture method. Crucially, models like colon organoids lessen the need for animal research, which raises ethical questions in addition to being expensive and difficult to grow and maintain. In this study, murine colon organoids derived from the colonic cryptic cells of C57BL/6 mice were employed to investigate the impact of Gum-Odina (GO)– modulated gut microbial metabolites on epithelial integrity in a colitis-like environment. This part of our study is the continuum of our previous study ^**[11]**^, where we proved that GO can significantly reduce the pathologies associated with DSS-induced colitis, including restoration of gut epithelial architecture and reduced pro-inflammatory cytokines TNF-_α_, IL-6, IFN-_γ_, IL-12, and induced anti-inflammatory cytokine IL-10. GO also proved to enhance gut microbiome diversity by stimulating the growth of the Firmicutes and Bacteroides bacteria, which competitively inhibit the growth of harmful gut bacteria. ^**[11]**^. GO also enhances the gut microbial metabolites, which are predominantly butyric acid, and other SCFAs including lactic acid, acetic acid and propionic acid ^**[16]**^. Here, to advance those findings, we used a colon organoid model to specifically evaluate the GO-modulated microbial metabolites on the gut epithelial integrity, independent of the systemic immunological impacts.

Over the years, SCFAs have been studied not only on the gut inflammation and barrier disruption ^**[17]**,**[18]**,**[19]**,**[20]**^ but also their levels are directly associated with hyperglycaemia ^**[21]**,**[22]**^, colorectal cancer ^**[23]**^, obesity ^**[24]**^, and autoimmune disorders ^**[25]**^. This leads us to the question if GO metabolites have an efficacious role in barrier enhancement in the IBD model. To address this question, we found the colon organoid model to be the best fit as it offers the real-time investigation of growth and morphology with the administration of GO metabolites over time. DSS has been proven to be the most common and valuable method which is vastly used to induce colitis both in vivo and in vitro ^**[26]**^. Disrupted epithelial border with increased cellular permeability has been the marker of DSS-induced colitis. Epithelial permeability is regulated by the tight junctional proteins which are the transmembrane proteins located at the basolateral side of the epithelial cells and responsible for maintaining cellular integrity. Among tight junction proteins, some are under the category of sealing protein which includes ZO-1, Occludin, and Claudin-1,4,5 whereas Claudin-2 is categorised as the pore-forming protein. The downregulation of Sealing proteins was observed in organoids grown with colitis metabolites however, organoids grown in the presence of GO metabolites showed enhanced expression of sealing proteins. This indicates GO metabolites are capable of inducing the tight junction protein synthesis in the epithelial cells which therefore can reduce the paracellular permeability of the intestine in the colitis condition. This finding corroborates with the study by Beisner et al where they have used inulin (prebiotic) and butyric acid (SCFA) to repair the disrupted gut barrier by increasing the expression of Occludin and ZO-1 in a western-style diet-induced obese mice model ^**[27]**^.

## Conclusion

In conclusion, this study highlights the protective role of Gum-Odina-derived gut microbial metabolites in restoring intestinal barrier integrity during colitis-associated conditions. Utilizing a murine colon organoid model, it was demonstrated that GO supplementation’s metabolites enhance the expression of sealing tight junction proteins such as Claudin-1, Claudin-4, Claudin-5, and Occludin, while maintaining JAM-A and Claudin-2 levels. This results in improved epithelial cohesion and lumen formation, underscoring GO metabolites’ barrier-protective effects. The findings are consistent with past DSS-induced colitis models, confirming the relevance of the organoid system. Overall, this work suggests Gum-Odina as a promising prebiotic therapy for improving gut permeability in colitis and has potential for future exploration in human-derived organoids for the inflammatory bowel disease therapies.

## Abbreviations

IBD: Inflammatory Bowel Disease
C-M: Colitis Metabolites (from SHIME)
GO-M: Gum Odina Metabolites (from SHIME)
C-GO-M: Colitis + Gum Odina Metabolites (treatment group in organoids)
SHIME: Simulator of Human Intestinal Microbial Ecosystem
3D: Three-dimensional
2D: Two-dimensional
PBS: Phosphate-Buffered Saline
BSA: Bovine Serum Albumin
DMEM/F-12: Dulbecco’s Modified Eagle Medium/Nutrient Mixture F-12
PFA: Paraformaldehyde
LPS: Lipopolysaccharide
SEM: Standard Error of the Mean
ANOVA: Analysis of Variance
qPCR: Quantitative (Real-Time) Polymerase Chain Reaction
FITC: Fluorescein Isothiocyanate
GO: Gum Odina
TJ: Tight Junction
ZO-1: Zonula Occludens-1
JAM-A: Junctional Adhesion Molecule-A
Cldn-1, -2, -4, -5*: Genes for Claudin-1, Claudin-2, Claudin-4, Claudin-5
Ocln: Gene for Occludin
Tjp1: Gene for Tight Junction Protein 1 (ZO-1)
SCFAs: Short-Chain Fatty Acids
DSS: Dextran Sodium Sulfate
GAPDH: Glyceraldehyde 3-Phosphate Dehydrogenase (housekeeping gene)
TNF-_α_: Tumor Necrosis Factor-alpha
IL-6: Interleukin-6
IFN-_γ_: Interferon-gamma
IL-12: Interleukin-12
IL-10: Interleukin-10

## Acknowledgements

The authors gratefully acknowledge the support of the Director of ICMR-National Institute for Research in Bacterial Infections (NIRBI), Kolkata, for facilitating this research. We extend our sincere thanks to Mr. Narayan Chandra Mondal for his diligent care of the mice. We acknowledge Animesh Gope for the technical assistance and use of the central instrumentation facilities. S.S. was supported by a CSIR fellowship during the course of this work.

## Author Contributions

Sohini Sikdar: Conceptualization, Methodology, Investigation, Formal analysis, Writing – Original Draft, Visualization.

Paulami Dutta: Methodology, Investigation, Formal analysis.

Debmalya Mitra: Methodology, Resources, Writing – Review & Editing, Supervision.

Amalesh Samanta: Lab space, equipment, and core infrastructure.

## Declaration of Generative AI and AI-Assisted Technologies in the Writing Process

During the preparation of this manuscript, the authors used ChatGPT and QuillBot to assist with grammatical corrections and to improve language clarity. All scientific content, data interpretation, results, and conclusions are entirely the authors’ own. After using these tools, the authors reviewed, edited, and took full responsibility for the final content of this work.

